# Gap-free telomere-to-telomere haplotypes assembly of the hybrid grouper ‘Yushuban’ (*Cromileptes altivelis* ♀ × *Epinephelus fuscoguttatus* ♂)

**DOI:** 10.64898/2026.09.20.752973

**Authors:** Youfeng Cai, Chongwei Wang, Ming Li, Yang Yang, Chunyou Cai, Sheng Lu, Songlin Chen

## Abstract

The hybrid grouper ‘Yushuban’ (*Cromileptes altivelis* ♀ × *Epinephelus fuscoguttatus* ♂) has shown promising performance in aquaculture trials, yet no complete reference genome has been available for this hybrid. We generated two gapless, telomere-to-telomere (T2T) haplotype-resolved genome assemblies of a single individual by integrating short-read, PacBio HiFi, Oxford Nanopore ultralong-read and Hi-C data. The maternal (Yushu_Calti_1.0) and paternal (Yushu_Efusc_1.0) haplotypes are 1,077.33 Mb and 1,083.36 Mb in length, with N50 values of 45.46 Mb and 45.55 Mb, respectively. Across all platforms, over 99.4% of reads mapped to each assembly, and Merqury-based quality values reached 57.7413 and 59.2068 (PacBio). BUSCO recovered 99.78% and 99.75% of complete core genes. We predicted 27,248 and 27,566 protein-coding genes, of which 98.52% and 99.23% were functionally annotated, and annotated telomeric motifs and repetitive elements (46.12% and 47.18% of each genome). These assemblies fill a gap in grouper genomic resources and provide a foundation for genetic dissection, parental genome comparison and molecular breeding of this hybrid.

## Background & Summary

Groupers (family *Epinephelidae*) are reef-associated fishes widely distributed in the tropical and subtropical waters of the Pacific and Indian Oceans and represent an economically important group of marine aquaculture species. The brown-marbled grouper (*Epinephelus fuscoguttatus*), which belongs to the genus *Epinephelus*, can spawn and milt throughout the year under captive conditions, making it an ideal parent for hybrid breeding^1^. The humpback grouper (*Cromileptes altivelis*) is the only species in the genus *Cromileptes*; although prized by consumers for its delicate flesh and attractive body shape, it grows slowly^2^, which has severely limited the expansion of its aquaculture industry. Hybridization has long been one of the most practical and widely adopted strategies in grouper breeding. Accumulating empirical evidence demonstrates that hybrid groupers can exhibit superior performance relative to their parental lines in growth rate and stress tolerance, as exemplified by the Hulong^3,4^ and Jinhu^5^ hybrid groupers. To address this problem, Hainan Chenhai Group crossed humpback grouper females with brown-marbled grouper males, successfully producing the hybrid grouper ‘Yushuban’ (*Cromileptes altivelis* ♀ × *Epinephelus fuscoguttatus* ♂), which showed promising performance in preliminary aquaculture trials.

Genome sequence largely determines the biological architecture of an organism, and a complete and accurate genome assembly is essential for a comprehensive understanding of organismal genetics and evolution^6^. Since the Telomere-to-Telomere (T2T) Consortium released the first complete, gap-free human genome (T2T-CHM13) in 2022^7^, genomics has entered an era in which complete, gap-free genomes are attainable. To date, genome assemblies have been reported for 24 grouper taxa (Epinephelinae), including 23 pure species and one hybrid, corresponding to 40 GenBank assembly records. Among them, chromosome-level or higher assemblies are available for 20 taxa, such as red-spotted grouper (*E. akaara*)^8^, leopard coral grouper (*Plectropomus leopardus*)^9^, kelp grouper (*E. moara*)^10^, potato grouper (*E. tukula*)^11^, brown-marbled grouper^12^, humpback grouper^13,14^, orange-spotted grouper (*E. coioides*)^15^ and Shanhu grouper (*E. fuscoguttatus* ♀ × *E. polyphekadion* ♂)^16^. However, only four pure grouper species have so far been assembled to the complete, gap-free T2T level: the giant grouper (*Epinephelus lanceolatus*)^17^, tomato hind (*Cephalopholis sonnerati*)^18^, blacktip grouper (*E. fasciatus*)^19^ and leopard coral grouper (*Plectropomus leopardus*)^20^. Most chromosome-level grouper assemblies still contain large gaps in repetitive regions, and their telomeres and centromeres often lack explicit annotation, which constrains high-resolution analyses of chromosomal evolution and structural variation. Generating a complete T2T assembly for Yushuban is therefore essential to fully resolve its dual-parental genomic architecture and support its genetic improvement.

In this study, we integrated next-generation sequencing (NGS), Pacific Biosciences (PacBio) HiFi sequencing, Oxford Nanopore Technology (ONT) ultralong-read sequencing and Hi-C data to construct high-quality T2T genomes of the hybrid grouper Yushuban. We successfully obtained two gap-free T2T haplotype-resolved genome assemblies (Fig. 1). Compared with the existing genome assemblies of its parental species (GCA_011397635.1^12^ and GCA_019925165.1^14^) (Table 3), the contiguity and completeness of all chromosomes were markedly improved. These two assemblies will provide important resources for population genetic studies, evolutionary analyses and the development of molecular breeding techniques for this hybrid and its parental species.

**Fig. 1.**
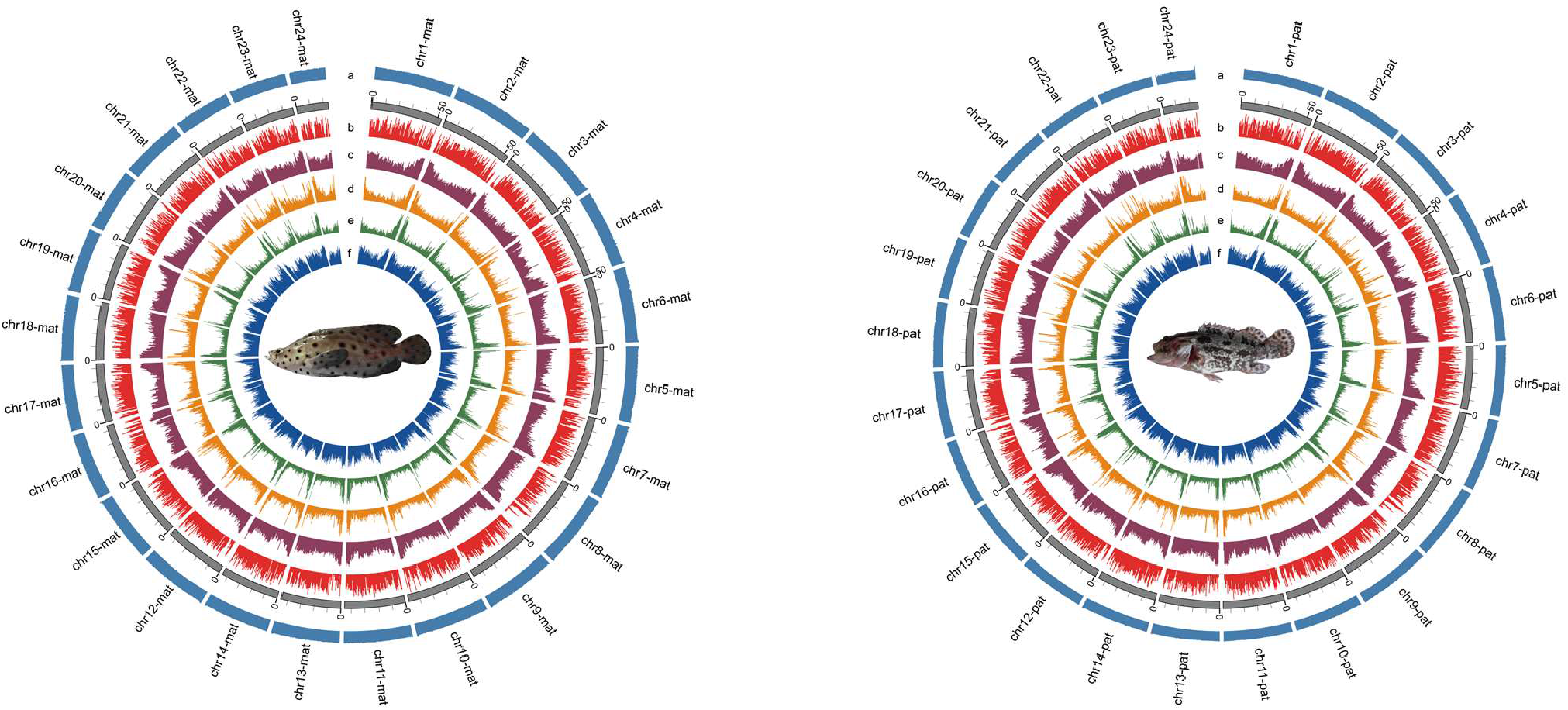
Circos plot illustrating the genome of *C. altivelis ♀* (left) *× E. fuscoguttatus ♂* (right). The plot includes the following components, arranged from outside to inside: (a) GC content distribution; (b) Gene density; (c) Density of total repetitive sequences; (d) LTR density; (e) LINE density; (f) DNA transposon (DNA-TE) density. The analysis was conducted using 500-kb genomic windows.

## Methods

### Ethics statement

All experimental procedures and sample collection methods in this study were approved by the Institutional Animal Care and Use Committee (IACUC) of the Yellow Sea Fisheries Research Institute, Chinese Academy of Fishery Sciences (CAFS) under approval No. YSFRI-2025017, and were performed in accordance with the Guidelines for the Care and Use of Laboratory Animals in China.

### Sample collection, library preparation and sequencing

A six-month-old individual of the hybrid grouper Yushuban (*C. altivelis ♀ × E. fuscoguttatus ♂*) was obtained from Lingshui Chenhai Seed Industry Co., Ltd., Hainan Province, China. After anaesthesia with MS-222, multiple tissues (muscle, fin, liver, blood, kidney, spleen, brain and heart) were collected and snap-frozen in liquid nitrogen to preserve the integrity of the genome and transcriptome. Muscle was used for whole-genome resequencing; muscle, blood and liver were used for PacBio HiFi sequencing, ONT ultralong-read sequencing and Hi-C sequencing, respectively; and total RNA was extracted from kidney, spleen, brain, fin, heart and muscle for transcriptome sequencing and genome annotation. DNA for second-generation sequencing was extracted and purified using an SDS-based magnetic-bead method, DNA for third-generation sequencing was extracted using SDS extraction with spin-column purification, and RNA was extracted using TRIzol. Sample quality was assessed with a NanoOne spectrophotometer (Yooning, China), a Qubit Flex Fluorometer (Thermo Fisher Scientific, USA) and agarose gel electrophoresis. RNA concentration was required to be ≥ 1 μg/μL with an A260/A280 ratio of 1.8-2.1 (the criterion for pure RNA). The RNA integrity number (RIN) was measured with an Agilent 2100 Bioanalyzer using the RNA Nano 6000 Assay Kit (Agilent Technologies, CA, USA), and only RNA with RIN ≥ 7.0 was retained to avoid degradation.

For NGS, sequencing libraries were constructed with the VAHTS Universal Plus DNA Library Prep Kit for MGI V2 following the standard operating procedure, and qualified libraries were sequenced on the DNBSEQ-T7RS platform (MGI Tech Co., Ltd., China) using the PE150 (150-bp paired-end) strategy. For HiFi sequencing, PacBio SMRTbell HiFi technology was used: libraries of 10 kb or 15 kb were constructed with the SMRTbell Express Template Prep Kit 2.0 and sequenced on the PacBio Revio platform, and circular consensus sequencing (CCS) mode was used to generate HiFi reads with accuracy ≥ Q30 (99.9%). For ONT ultralong sequencing, nanopore libraries were prepared with the ONT SQK-LSK114 kit and loaded onto R10.4.1 flow cells, followed by sequencing for 48-72 h on a PromethION instrument (Oxford Nanopore Technologies, UK). For Hi-C, chromatin conformation was fixed with formaldehyde, cells were lysed and digested with DpnII to generate sticky ends, and the DNA ends were end-repaired with concomitant incorporation of biotin-14-dATP. DNA fragments were ligated with DNA ligase, and cross-links were released by proteinase digestion. The DNA was purified, randomly sheared into 300–500-bp fragments, and biotin-labelled fragments were captured on streptavidin magnetic beads. The fragments were then end-repaired, A-tailed and ligated to sequencing adapters, and the appropriate number of PCR amplification cycles was determined before final purification. After library quality control, sequencing was performed on the DNBSEQ-T7RS platform (MGI Tech Co., Ltd., China). In total, 101.68 Gb of NGS data, 147.52 Gb of PacBio HiFi data, 168.63 Gb of ONT ultralong data and 226.51 Gb of Hi-C data were generated for downstream analysis (Table 1).

**Table 1.**
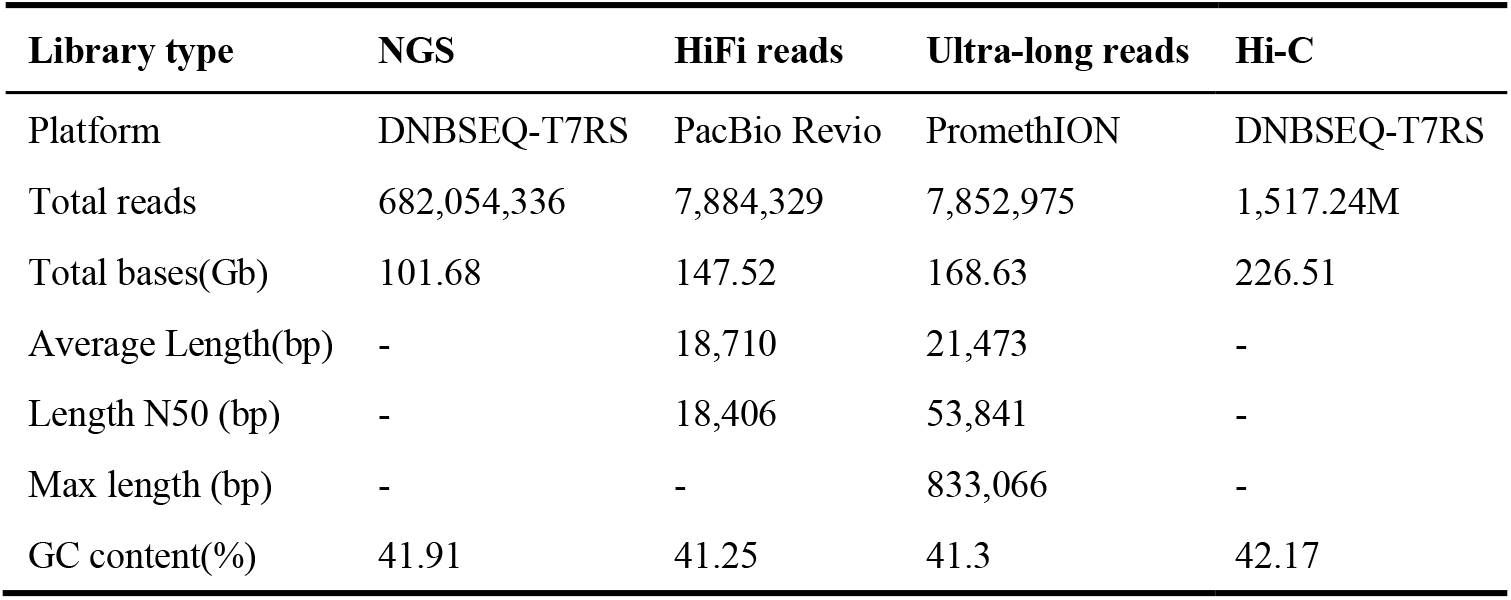
Statistics of sequencing data for genome assembly.

| Library type | NGS | HiFi reads | Ultra-long reads | Hi-C |
| --- | --- | --- | --- | --- |
| Platform | DNBSEQ-T7RS | PacBio Revio | PromethION | DNBSEQ-T7RS |
| Total reads | 682,054,336 | 7,884,329 | 7,852,975 | 1,517.24M |
| Total bases(Gb) | 101.68 | 147.52 | 168.63 | 226.51 |
| Average Length(bp) | - | 18,710 | 21,473 | - |
| Length N50 (bp) | - | 18,406 | 53,841 | - |
| Max length (bp) | - | - | 833,066 | - |
| GC content(%) | 41.91 | 41.25 | 41.3 | 42.17 |

### Genome survey

Reads were quality-controlled with fastp^21^ (v0.23.4, parameter: -l 50 --dedup --dup_calc_accuracy 3), which trims reads containing adapter sequences, ambiguous (N) bases and low-quality bases; low-quality, too-short and PCR-duplicated reads were removed, and only valid read pairs were retained. Quality was assessed with FastQC^22^ (v0.12.1, default parameters). To detect sample contamination and verify data reliability, 20,000 randomly selected reads were aligned to the NCBI NT database with Blastn^23^. K-mer frequency distributions were obtained with FastK^24^ (v1.1.0, parameter: -k 21); based on the k-mer (k = 21) frequencies, genome characteristics were estimated with GCE^25^ (v1.0.2, parameter: -k 21) and ploidy was analysed with Smudgeplot^26^ (v0.2.3dev, parameter: -k 21). In parallel, k-mer analysis was performed with Jellyfish^24^ (v2.3.0, parameter: -k 21) and genome characteristics were estimated with GenomeScope^26^ (v2.0, parameter: -k 21 -p 2). The k-mer analysis indicated a haploid genome size of approximately 1,018 Mb, a heterozygosity rate of 2.34-3.18% and a repeat content of 22.66-28.106% (Table 2).

**Table 2.** Genome survey of *C. altivelis ♀ × E. fuscoguttatus ♂*.

| Tool | K-mer size | K-mer number | K-mer depth(×) | Genome size(bp) | Heterozygous-ratio(%) | Repeat(%) |
| --- | --- | --- | --- | --- | --- | --- |
| GCE | 21 | 85,794,590,315 | 83 | 1,017,822,718 | 2.34 | 22.66 |
| Genomescope | 21 | - | - | 1,017,987,802 | 3.18 | 28.106 |

**Table 3.** Statistics of *C. altivelis ♀ × E. fuscoguttatus ♂* genome assemblies.

| Assembly | OUC_Cal_1.0<sup>14</sup> | *E.fuscoguttatus*.final_Chr_v1<sup>12</sup> | Yushu_Calti_1.0 | Yushu_Efusc_1.0 |
| --- | --- | --- | --- | --- |
| species | *C. altivelis* | *E. fuscoguttatus* | *C. altivelis* | *E. fuscoguttatus* |
| Total genome length (bp) | 1,083,008,411 | 1,046,995,804 | 1,077,334,252 | 1,083,364,598 |
| Total chromosome length (bp) | 1,082,955,183 | 1,046,975,304 | 1,077,334,252 | 1,083,364,598 |
| No. chromosome | 24 | 24 | 24 | 24 |
| No. scaffolds | 306 | 25 | 24 | 24 |
| Scaffold N50 (bp) | 43,784,314 | 44,419,306 | 45,462,755 | 45,552,130 |
| No. contigs | 824 | 230 | 24 | 24 |
| Contig N50 (bp) | 5,160,563 | 13,857,438 | 45,462,755 | 45,552,130 |
| GC percent (%) | 41.5 | 41.5 | 41.24 | 41.36 |
| Complete BUSCOs (Proteins/%) | 3,263/97.3 | 3,625/99.59 | 7,191/99.78 | 7,189/99.75 |
| Complete Single-Copy BUSCOs (Proteins/%) | 3,230/96.3 | 3,598/98.85 | 7,170/99.49 | 7,168/99.46 |
| Complete Duplicated BUSCOs (Proteins/%) | 33/1.0 | 27/0.74 | 21/0.29 | 21/0.29 |
| Fragmented BUSCOs (Proteins/%) | 47/1.4 | 8/0.22 | 1/0.01 | 2/0.03 |
| Missing BUSCOs (Proteins/%) | 44/1.3 | 7/0.19 | 15/0.21 | 16/0.22 |

### Genome assembly and gap filling

The initial assembly was performed with hifiasm^27^ (v0.25.0-r726, parameter: --n-hap 2) using the PacBio HiFi, ONT ultralong and Hi-C reads. For this highly heterozygous genome, Purge Haplotigs^28^ (v1.1.3, default parameters) was used to remove redundant haplotypic contigs based on sequencing depth and sequence similarity. The assembly was then aligned to the NCBI NT database (https://ftp.ncbi.nlm.nih.gov/blast/db/) with Blastn^23^ (v2.11.0+, parameter: -evalue 0.00001 -max_hsps 1), and organellar and exogenous contaminating sequences were identified and removed based on the alignment results, yielding a high-quality genome assembly. Hi-C data were pre-processed and quality-controlled with fastp^21^ (v0.23.4, parameter: -l 50 --dedup --dup_calc_accuracy 3), assessed with FastQC^22^ (v0.12.1, default parameters), and the effective reads were evaluated with HiCUP^29^ (v0.7.2, parameter: --NM 3) to verify Hi-C library quality and data validity. Clean reads were aligned to the genome with Juicer^30^ (v1.6, default parameters), and the alignments were used by 3D-DNA^31^ and HapHiC^32^ (v1.0.2, parameter: --NM 3) for contig clustering, ordering and orienting; the higher-quality version was selected. The assembly was then manually refined with JuiceBox^33^ (v1.11.08, default parameters), ultimately producing chromosome-level genome assemblies.

Initial gap filling was performed with TGS-GapCloser^34^ (v1.2.0, parameter: --min_nread 10), which uses the coverage relationship between Nanopore ultralong reads and the assembled contigs to fill inter-contig gaps and extend contigs. After gap filling, reads were re-aligned to the updated genome, and the read distribution around the updated sites was inspected to verify the accuracy of the filling. In parallel, gap filling and telomere extension were performed by local assembly of the gap and telomere regions. The results of the two approaches were integrated and the optimal method was used to update the genome based on the gap-filling and extension performance. Finally, the genome was polished with high-quality short-read and HiFi data. Telomere regions were identified by a genome-wide search for the telomeric repeat motif (TTAGGG)n, and telomeric motifs were counted within the terminal 100-kb window of each chromosome (at least four copies of the motif were required). Next-generation resequencing data from fin-clip samples of non-parental individuals of the two parental species (*C. altivelis* and *E. fuscoguttatus*) were mapped to the merged hap1/hap2 genome assembly, and read-depth-based genotyping was performed to assign the two haplotype sets to their respective parental species.(Fig. S1; Table S1) Ultimately, we obtained two gapless T2T haplotype assemblies of the hybrid grouper Yushuban, maternal (Yushu_Calti_1.0) and paternal (Yushu_Efusc_1.0), of 1,077.33 Mb (N50, 45.46 Mb) and 1,083.36 Mb (N50, 45.55 Mb)(Fig. 1, 2, 3; Table 4), respectively. Compared with the published reference genomes (OUC_Cal_1.0)^14^ and (*E. fuscoguttatus*.final_Chr_v1)^12^, both assemblies showed markedly improved completeness and contiguity (Table 3). Synteny analysis between the two haplotypes identified 135 syntenic blocks containing a total of 23,895 gene pairs, with an average of 177 gene pairs per block (Fig. S2).

**Table 4.** Summary of chromosomal assembly features, including length and telomere repeat units.

| Yushu_Calti_1.0 |  |  |  | Yushu_Efusc_1.0 |  |  |  |
| --- | --- | --- | --- | --- | --- | --- | --- |
| #chr | Length(bp) | Number of start repeat unit | Number of end repeat unit | #chr | Length(bp) | Number of start repeat unit | Number of end repeat unit |
| chr1-mat | 53,082,293 | 0 | 414 | chr1-pat | 53,859,551 | 745 | 677 |
| chr2-mat | 53,506,655 | 904 | 406 | chr2-pat | 53,571,245 | 1283 | 156 |
| chr3-mat | 51,331,874 | 1069 | 0 | chr3-pat | 51,359,725 | 864 | 494 |
| chr4-mat | 50,206,427 | 741 | 0 | chr4-pat | 49,436,801 | 574 | 1182 |
| chr5-mat | 49,831,582 | 570 | 0 | chr5-pat | 47,269,310 | 0 | 313 |
| chr6-mat | 49,470,014 | 324 | 1600 | chr6-pat | 49,829,303 | 506 | 580 |
| chr7-mat | 49,421,979 | 445 | 789 | chr7-pat | 48,273,949 | 1262 | 0 |
| chr8-mat | 48,669,027 | 719 | 750 | chr8-pat | 48,900,896 | 536 | 744 |
| chr9-mat | 47,490,068 | 0 | 0 | chr9-pat | 48,601,480 | 0 | 632 |
| chr10-mat | 48,467,462 | 307 | 494 | chr10-pat | 46,115,256 | 201 | 0 |
| chr11-mat | 45,014,776 | 417 | 859 | chr11-pat | 45,552,130 | 149 | 0 |
| chr12-mat | 45,462,755 | 957 | 432 | chr12-pat | 44,560,888 | 585 | 757 |
| chr13-mat | 44,826,401 | 634 | 434 | chr13-pat | 45,405,903 | 0 | 742 |
| chr14-mat | 44,258,976 | 579 | 361 | chr14-pat | 45,367,845 | 0 | 652 |
| chr15-mat | 44,298,748 | 603 | 1079 | chr15-pat | 45,068,635 | 499 | 1351 |
| chr16-mat | 43,871,399 | 596 | 680 | chr16-pat | 45,010,519 | 0 | 0 |
| chr17-mat | 43,429,929 | 637 | 1271 | chr17-pat | 44,514,296 | 636 | 1231 |
| chr18-mat | 43,174,569 | 0 | 566 | chr18-pat | 43,854,524 | 0 | 705 |
| chr19-mat | 42,403,492 | 627 | 0 | chr19-pat | 41,384,776 | 1173 | 415 |
| chr20-mat | 40,330,632 | 374 | 515 | chr20-pat | 40,858,756 | 553 | 158 |
| chr21-mat | 40,844,046 | 349 | 995 | chr21-pat | 40,371,890 | 448 | 642 |
| chr22-mat | 36,588,708 | 698 | 645 | chr22-pat | 40,544,870 | 1015 | 977 |
| chr23-mat | 37,472,238 | 1126 | 0 | chr23-pat | 37,551,854 | 207 | 0 |
| chr24-mat | 23,880,202 | 31 | 0 | chr24-pat | 26,100,196 | 157 | 966 |
| total | 1,077,334,252 |  |  | total | 1,083,364,598 |  |  |

**Fig. 2.**
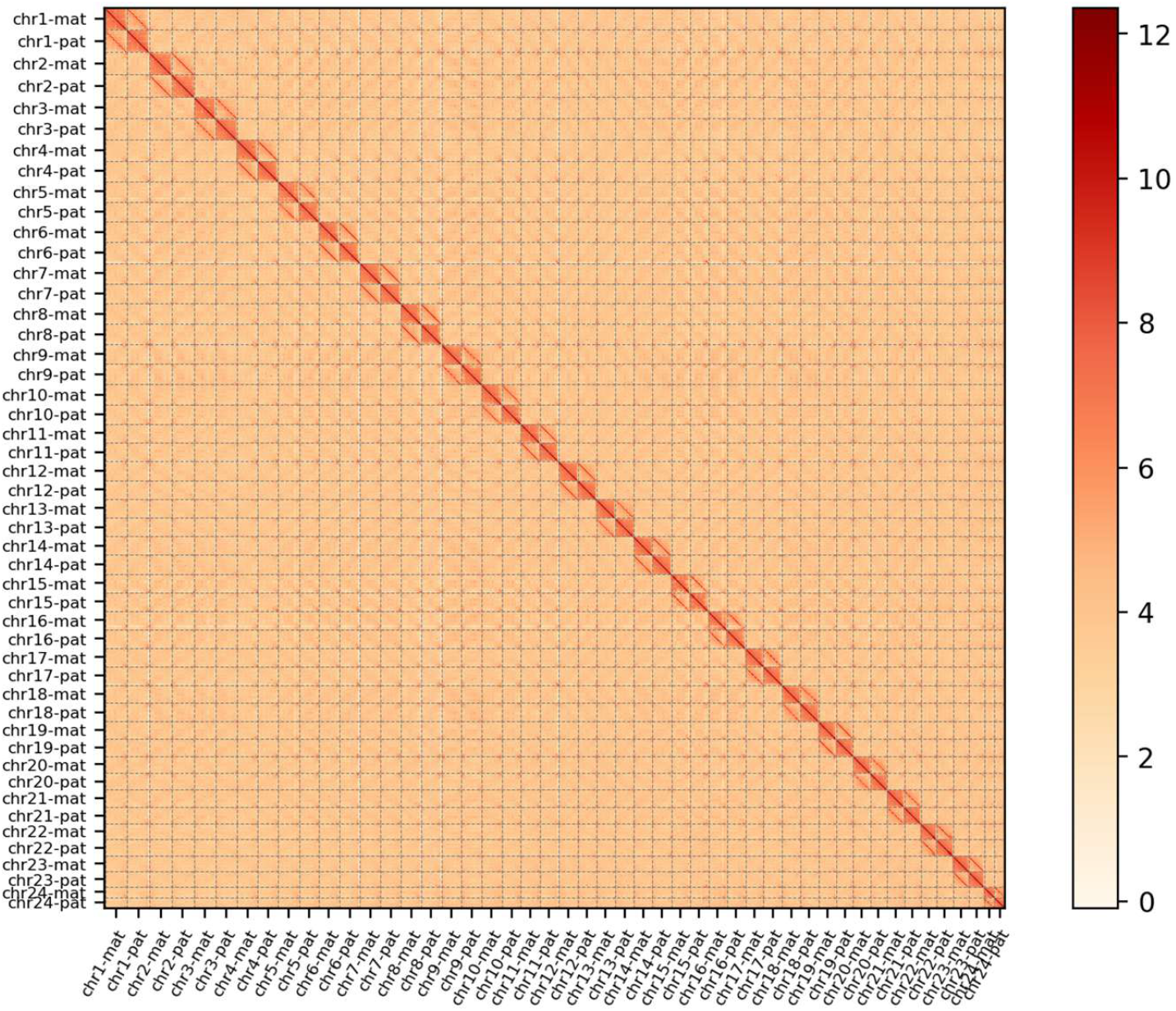
Hi-C contact map of two haplotype genome assemblies at a bin size of 1 Mb, showing clear chromosome-scale scaffolding and interaction signals.

**Fig. 3.**
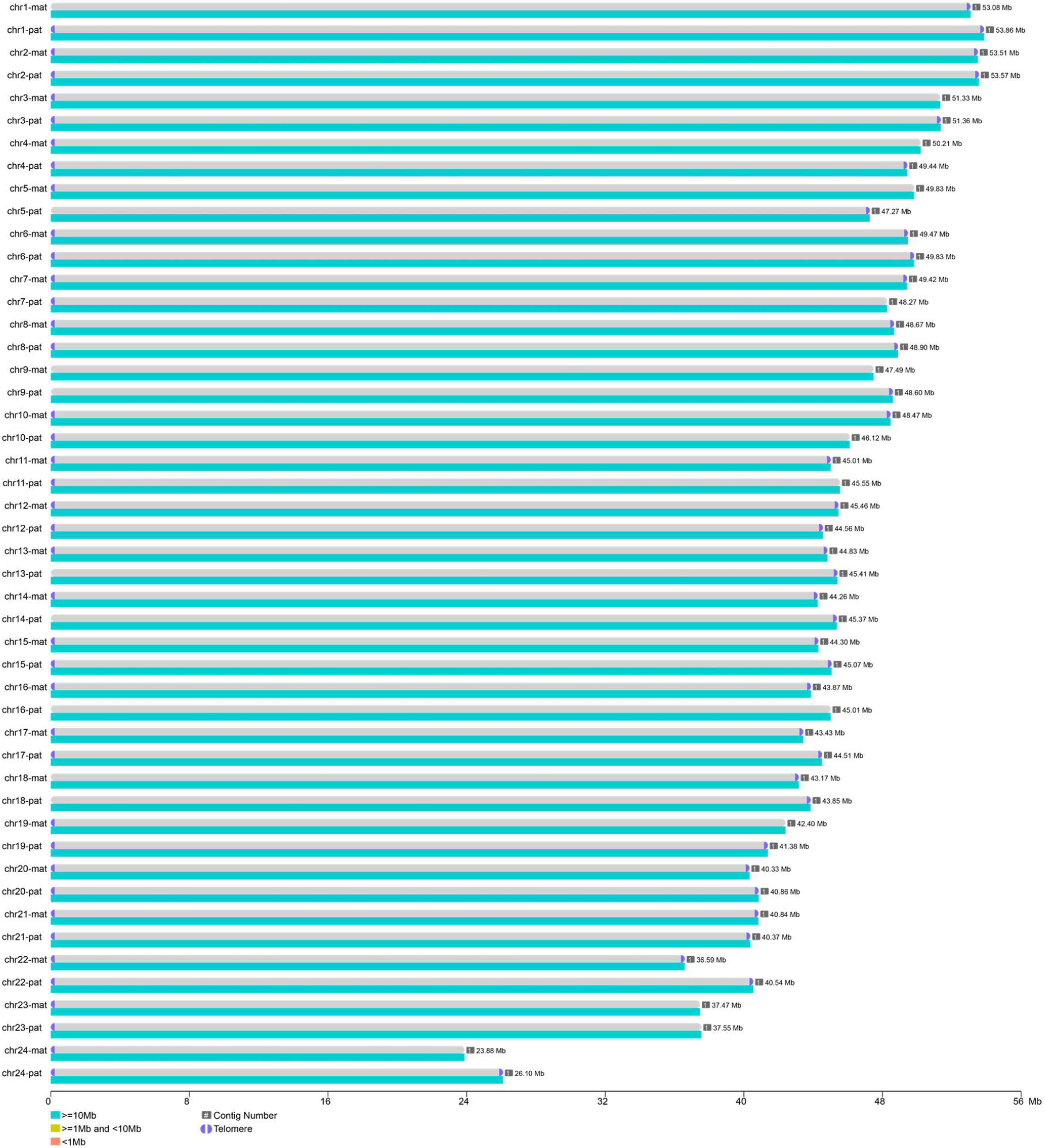
Contig distribution on chromosomes in the genome. The blue sections at each end of chromosomes represent the identified telomeres.

### Repeat sequence annotation

Repetitive elements were annotated by combining de novo prediction, hom ology-based search and an integrated strategy. TRF^35^ (v4.10.0-rc.2, parameter: 2 7 7 80 10 50 2000 -d -h -m) was used for de novo prediction of tandem repeats based on the sequence features. RepeatMasker^36^ (v4.2.1, parameter: -nolow -no_is -norna) and its built-in tool RepeatProteinMask^36^ (v4.2.1, default parameters) were used to search the combined database (https://www.girinst.org/server/RepBase/protected/repeatmaskerlibraries/) for the homology-based identification of known repeats. RepeatModeler^37^ (v2.0.7, default parameters) a nd LTR_FINDER_parallel^38^ (v1.1, parameter: -harvest_out -size 1000000 -time 300) were combined to construct a species-specific repeat library, which was then used as a reference set to re-run RepeatMasker for comprehensive annotation, improving the sensitivity and accuracy of the prediction. In total, 496.83 Mb and 5 11.12 Mb of repetitive sequences were identified in Yushu_Calti_1.0 and Yushu_Efusc_1.0, accounting for 46.12% and 47.18% of the respective genomes. DNA transposons accounted for 23.08% and 23.89% of the genomes, long terminal repeats (LTRs) for 4.36% and 5.24%, long interspersed nuclear elements (LINEs) fo r 7.6% and 7.07%, and short interspersed nuclear elements (SINEs) for 0.98% and 1.09% (Fig. 1; Tables 5 a nd 6).

**Table 5.** Statistics of repeat sequence annotation.

| Type | Yushu_Calti_1.0 |  | Yushu_Efusc_1.0 |  |
| --- | --- | --- | --- | --- |
|  | Length (Bp) | % in genome | Length (Bp) | % in genome |
| DNA | 248,610,117 | 23.08 | 258,803,174 | 23.89 |
| LINE | 81,905,828 | 7.6 | 76,618,501 | 7.07 |
| SINE | 10,592,307 | 0.98 | 11,779,569 | 1.09 |
| LTR | 46,926,755 | 4.36 | 56,750,356 | 5.24 |
| Satellite | 3,776,036 | 0.35 | 2,876,276 | 0.27 |
| Simple_repeat | 0 | 0 | 0 | 0 |
| Other | 35,881 | 0 | 40,586 | 0 |
| Unknown | 126,741,910 | 11.76 | 128,771,852 | 11.89 |
| Total | 496,830,139 | 46.12 | 511,117,787 | 47.18 |

**Table 6.**
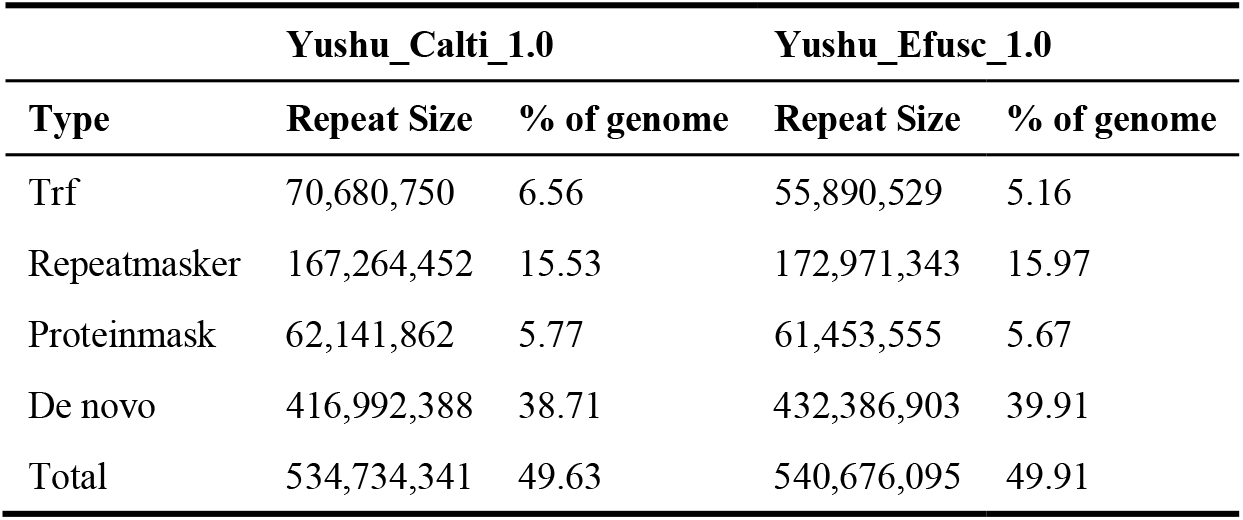
Summary of transposon element families based on various methods.

| Type | Yushu_Calti_1.0 |  | Yushu_Efusc_1.0 |  |
| --- | --- | --- | --- | --- |
|  | Repeat Size | % of genome | Repeat Size | % of genome |
| Trf | 70,680,750 | 6.56 | 55,890,529 | 5.16 |
| Repeatmasker | 167,264,452 | 15.53 | 172,971,343 | 15.97 |
| Proteinmask | 62,141,862 | 5.77 | 61,453,555 | 5.67 |
| De novo | 416,992,388 | 38.71 | 432,386,903 | 39.91 |
| Total | 534,734,341 | 49.63 | 540,676,095 | 49.91 |

### Protein-coding gene annotation

For homology-based prediction, protein sequences of closely related species were aligned to the target genomes with miniprot^39^ (v0.18, default parameters) and Liftoff^40^ (v1.6.3, parameter: -a 0.5 -s 0.5). Ab initio prediction was performed with AUGUSTUS^41^ (v3.3.2, default parameters) and Genscan^42^ using the repeat-masked genomes. For RNA-seq data, reads were aligned to the genome with HISAT2^43^ (v2.1.0, parameter: --dta), transcripts were assembled with StringTie^44^ (v1.3.5, default parameters), and coding regions were predicted from the transcripts with TransDecoder (v5.5.0, default parameters, https://github.com/TransDecoder/TransDecoder). For Iso-Seq data, transcripts were extracted with IsoSeq3 (v4.3.0, default parameters, https://github.com/PacificBiosciences/IsoSeq), aligned to the genome with minimap2^45^ (v2.24-r1122, parameter: -ax map-ont/hifi/pb), converted with spliced_bam2gff, and coding regions were predicted with TransDecoder (v5.5.0, default parameters). Gene models were also predicted with compleasm^46^ (v0.2.7, parameter: compleasm run -m busco) based on the BUSCO set. All the above evidence was integrated with MAKER2^47^ (v2.31.10, parameter: max_dna_len=3000000, min_contig=10000, pred_flank=500, min_protein=30) to obtain a non-redundant gene set, which was then refined with HiFAP (Wuhan OneMore, https://www.onemore tech.com) to produce a high-quality consensus gene set. Ultimately, 27,248 and 27,566 protein-coding genes were identified in the two assemblies (Tables 7 and 8), yielding 36,001 and 36,629 transcripts, respectively (Table 9). Compared with five closely related species, the gene structures were highly consistent across the four dimensions examined (Fig. 4).

**Table 7.**
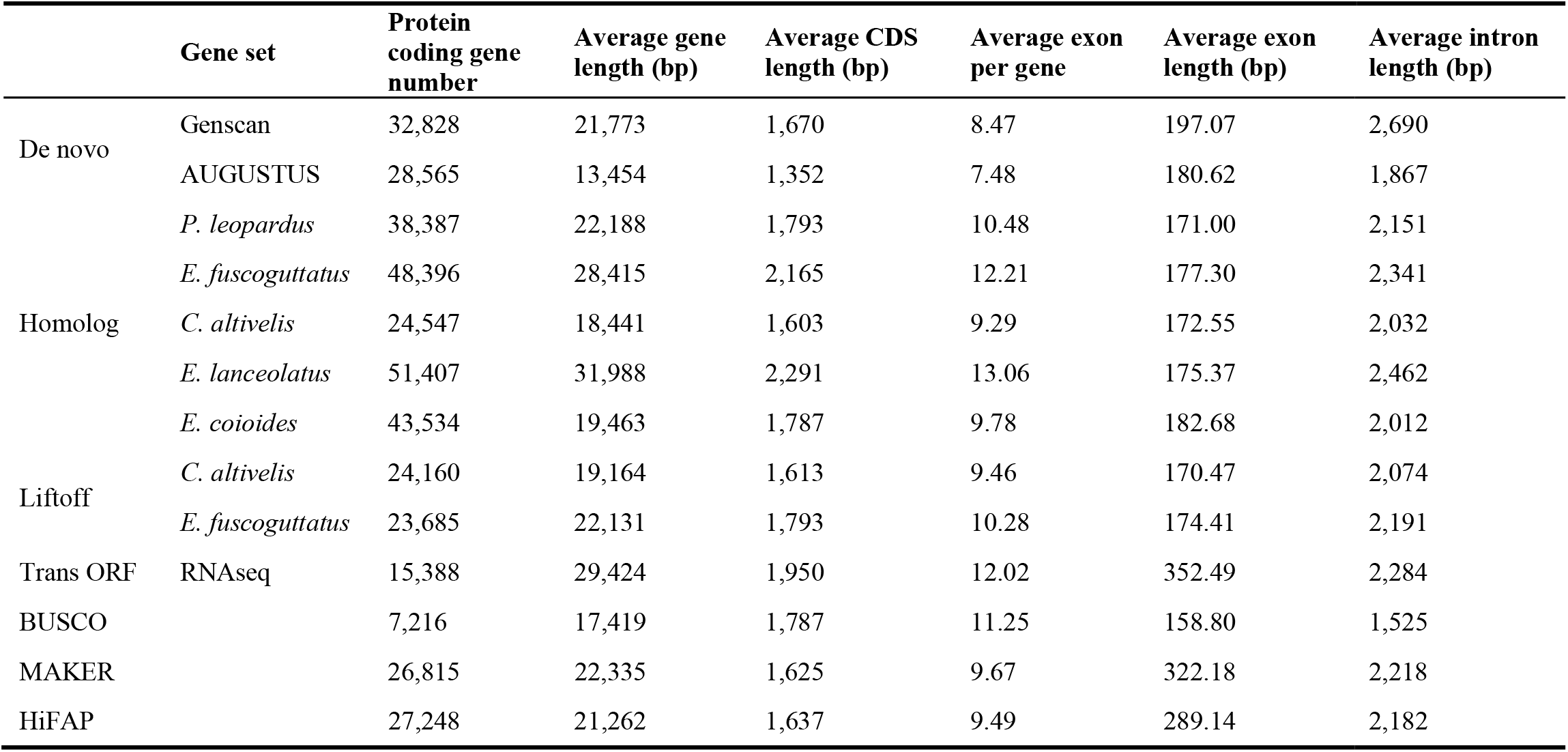
Gene prediction statistics of Yushu_Calti_1.0.

**Table 8.**
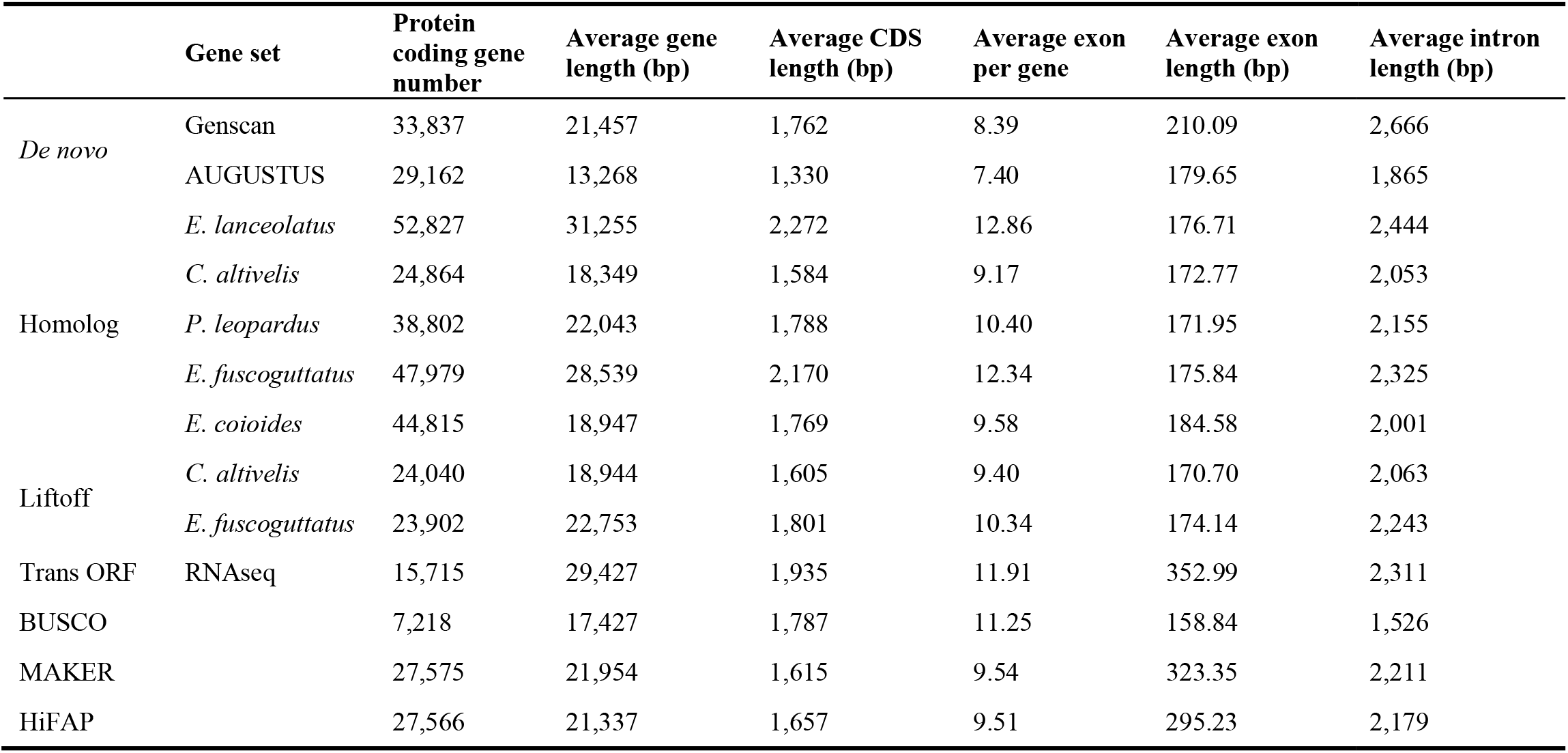
Gene prediction statistics of Yushu_Efusc_1.0.

**Table 9.** Statistics of gene sets with isoforms.

|  | Yushu_Calti_1.0 | Yushu_Efusc_1.0 |
| --- | --- | --- |
| Total gene number | 27,248 | 27,566 |
| Total transcript number | 36,001 | 36,629 |
| Mean length of transcripts (bp) | 24,683.77 | 25,110.94 |
| Median length of transcripts (bp) | 10,958 | 10,963 |
| Mean length of exon (bp) | 172.43 | 173.64 |
| Median length of exon (bp) | 124 | 125 |

**Fig. 4.**
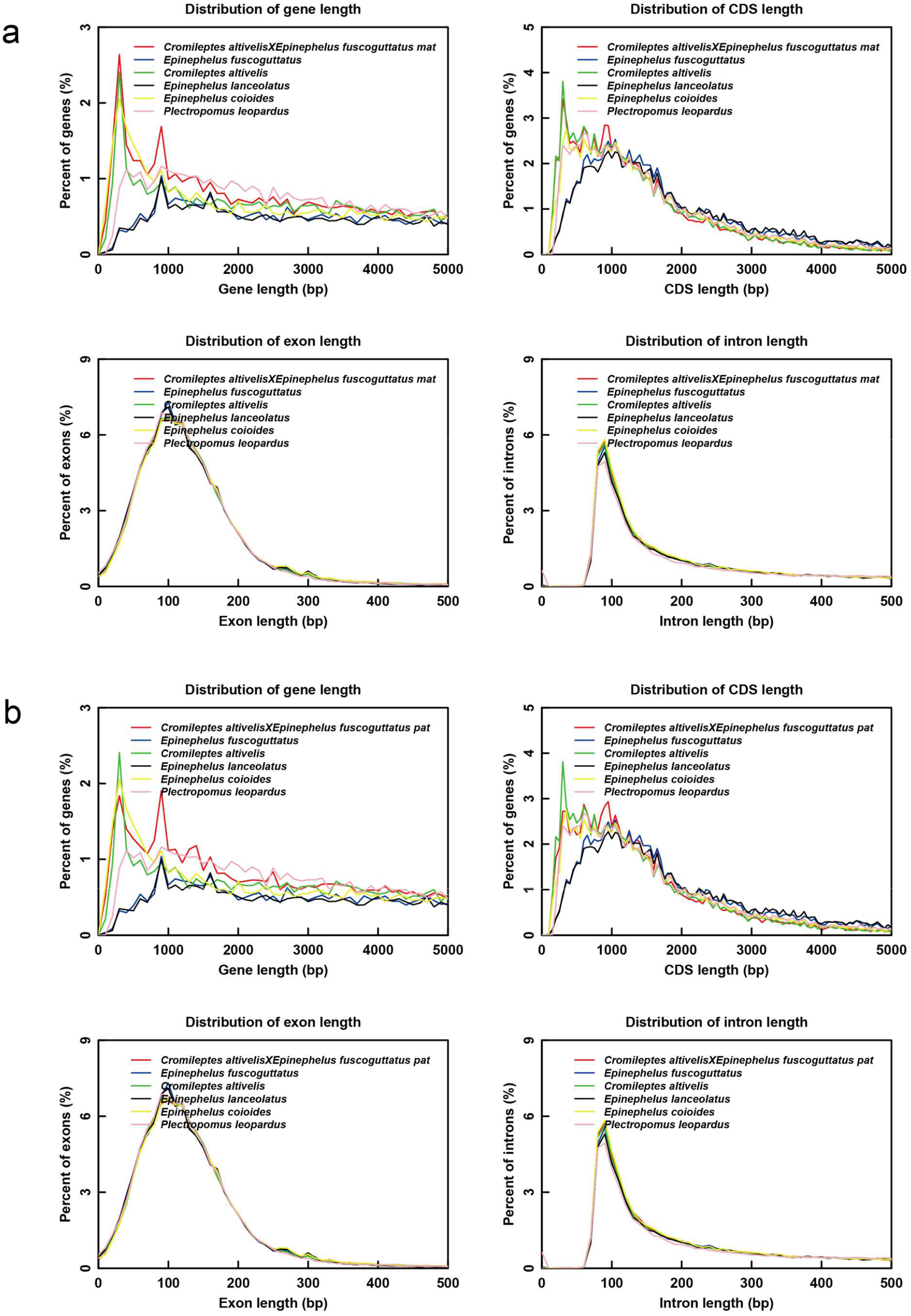
Distribution of gene length, CDS length, exon length and intron length among related species. (a) Yushu_Calti_1.0; (b) Yushu_Efusc_1.0.

### Gene function annotation

To characterise the functions and metabolic pathways of the predicted genes, systematic functional annotation of the encoded protein sequences was performed. Protein sequences were aligned with diamond^48^ (v2.1.11.165, parameter: blastp --max-hsps 1 --outfmt 6 -e 1e-5) against the NR^49^ (NCBI Non-redundant Protein Database), Swiss-Prot^50^, TrEMBL^50^, KOG^51^, REBASE^52^, CYPED^53^, TCDB^54^ and AnimalTFDB^55^ (v4.0) databases, and the highest-scoring hit was retained as the final annotation. KofamScan^56^ (v1.3.0, parameter: -E 1e-5 --format detail-tsv) was used to align protein sequences against the KOfam database (v117.0) for KEGG^57^ KO annotation, and the KEGG BRITE database was used to interpret metabolic pathways. InterPro^58^, GO^59^ and Pfam^60^ annotations were performed with InterProScan^61^ (v5.76-107.0, parameter: --seqtype p --formats TSV --goterms --pathways -dp). CAZy^62^ annotation was performed with dbCAN^62^ (v5.1.2, parameters: --e_value_threshold 1e-102, --coverage_threshold_dbcan 0.35, --e_value_threshold_dbcan 1e-15, --coverage_threshold_dbsub 0.35, --e_value_threshold_dbsub 1e-15). An E-value threshold of 1e-5 was used for all annotations to ensure reliable results. Transmembrane domains were annotated with TMHMM^63^ (v2.0c, default parameters), secreted proteins were predicted with SignalP^64^ (v6.0h, parameter: --mode slow-sequential), and protein subcellular localisation was predicted with WoLF PSORT^65^ (v0.2, default parameters). In total, 26,845 and 27,355 of the predicted genes in the two assemblies were annotated, corresponding to 98.52% and 99.23% of the gene sets (Table 10).

**Table 10.**
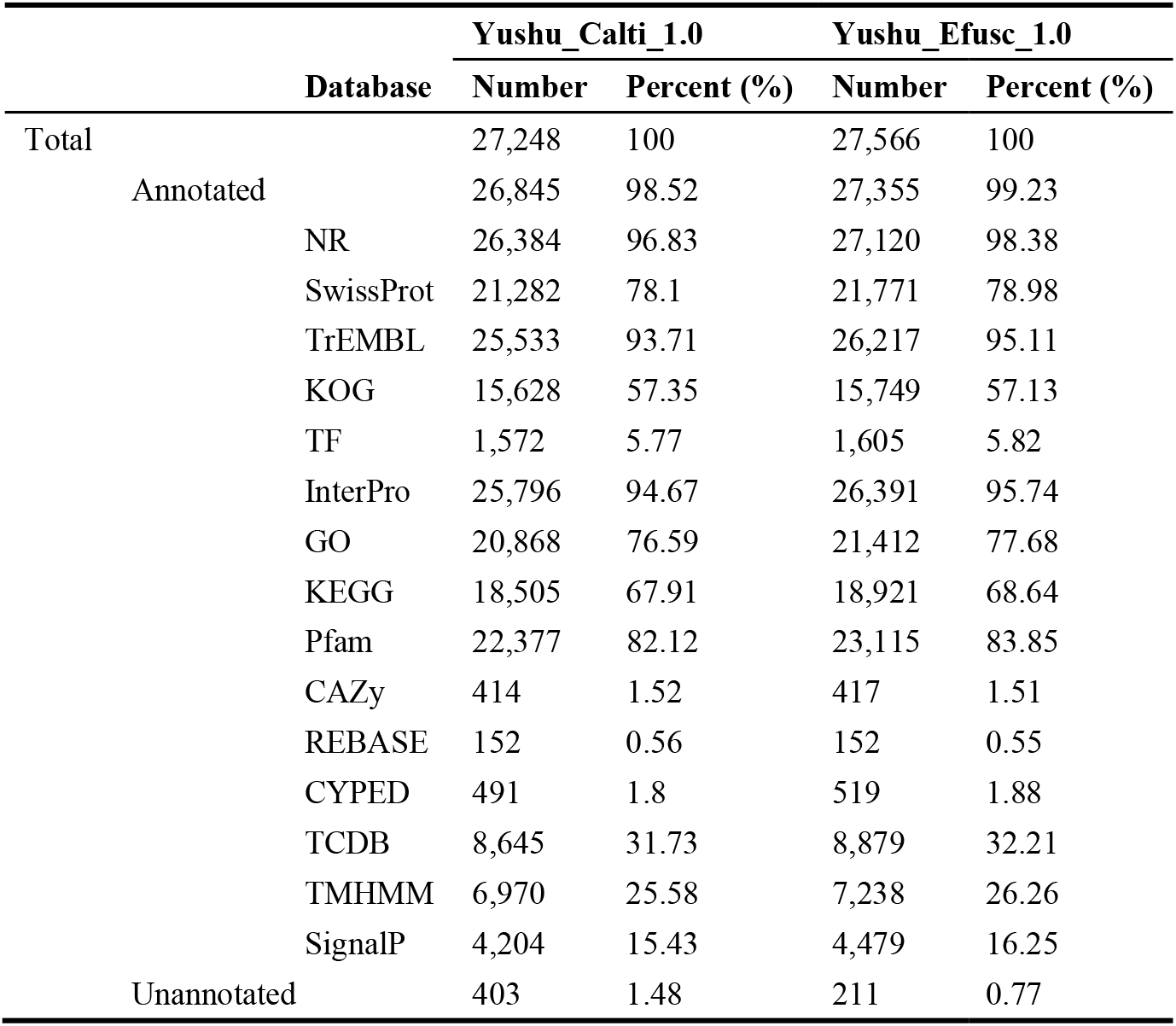
Statistics of functional annotation.

### Non-coding RNA annotation

Non-coding RNAs were annotated with the cmscan tool of Infernal^66^ (v1.1.5, default parameters) against the Rfam database (v14.1). rRNA genes were predicted by cross-validation with both barrnap (v1.10.5, default parameters, https://github.com/tseemann/barrnap) and cmscan, and tRNA genes were identified by precise structural and sequence recognition with tRNAscan-SE^67^ (v2.0.12, default parameters). The results are summarised in Table 11.

**Table 11.**
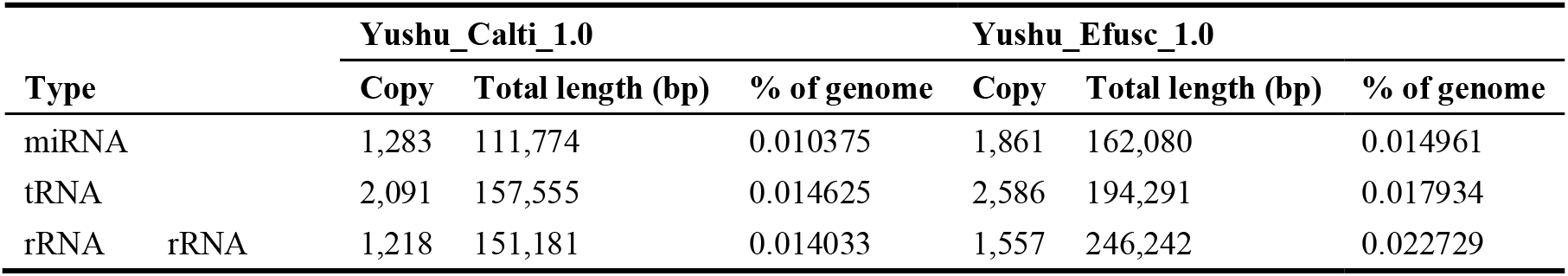
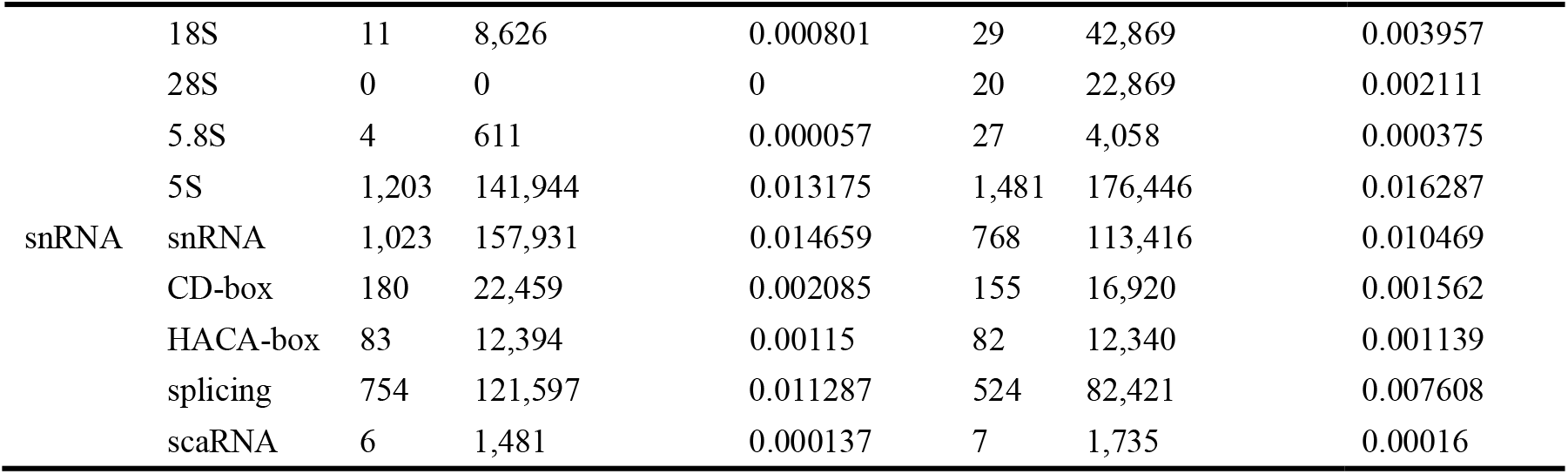
Statistics of non-coding RNA annotation.

## Data Records

The raw data (including NGS, PacBio Hi-Fi, ONT ultra-long, Hi-C, and RNA sequencing data) has been submitted to the NCBI Sequence Read Archive under BioProject number PRJNA1511585, with accession numbers SRR40183358^68^, SRR40183353^69^, SRR40183354^70^, SRR40183355^71^, and SRR40183352^72^. The two genome assemblies have been deposited into NCBI GenBank under the accession JCDUVQ000000000^73^ (Yushu_Calti_1.0) and JCDUVR000000000^74^ (Yushu_Efusc_1.0). The version described in this paper are version JCDUVQ010000000 and JCDUVR010000000. Additionally, the two genome assemblies and genome annotation files were submitted to FigShare^75^.

## Technical Validation

### Genome assembly and annotation evaluation

For contiguity, short reads, PacBio long reads and Nanopore ultralong reads were aligned to the genome with bwa^76^ (v0.7.12-r1039, parameter: -M) and minimap2^45^ (v2.24-r1122, parameter: -ax map-ont/hifi/pb), respectively. The mapping rates of two assemblies were more than 99.4%, and the coverage rates were more than 98.63% (at least 20X) (Tables 12). For completeness, BUSCO^77^ (v6.0.0 (odb12), parameter: --miniprot -m genome -f --offline) and Compleasm^46^ (v0.2.7 (odb12), parameter: compleasm run -m busco) were used to search the highly conserved orthologue sets in the OrthoDB database (https://busco-data.ezlab.org/v5/data/lineages/), and the numbers and proportions of complete (single-copy and duplicated), fragmented and missing BUSCOs were calculated. A total of 7,207 genes were searched, of which 7,191 (99.78%) and 7,189 (99.75%) were identified as complete, and 7,170 (99.49%) and 7,168 (99.46%) were single-copy and conserved, in the two assemblies. Only a small proportion of genes (at most 0.25%) were fragmented or missing (Table 13). For base accuracy, assembly quality values (QV) of two genomes were calculated with Merqury^78^ (v1.3, default parameters), yielding 46.295 and 46.0731 for the short-read data and 57.7413 and 59.2068 for the PacBio long-read data (Table 14).

**Table 12.** Statistics of alignment results.

| Data type | Yushu_Calti_1.0 |  |  | Yushu_Efusc_1.0 |  |  |
| --- | --- | --- | --- | --- | --- | --- |
|  | short-read | PacBio long-read | Nanopore ultra-long read | short-read | PacBio long-read | Nanopore ultra-long read |
| Mapping rate (%) | 99.42 | 99.99 | 99.59 | 99.5 | 99.98 | 99.62 |
| Paired mapping rate (%) | 94.46 | - | - | 94.99 | - | - |
| Average sequencing depth | 91.06 | 128.48 | 147.28 | 90.88 | 129.15 | 147.6 |
| Coverage (%) | 99.85 | 99.93 | 100 | 99.9 | 99.96 | 100 |
| Coverage at least 4X (%) | 99.67 | 99.87 | 99.99 | 99.75 | 99.91 | 100 |
| Coverage at least 10X (%) | 99.35 | 99.8 | 99.98 | 99.48 | 99.87 | 99.99 |
| Coverage at least 20X (%) | 98.63 | 99.69 | 99.97 | 98.86 | 99.81 | 99.98 |

**Table 13.** BUSCO assessment result.

|  | Yushu_Calti_1.0 |  |  |  | Yushu_Efusc_1.0 |  |  |  |
| --- | --- | --- | --- | --- | --- | --- | --- | --- |
|  | Assembly |  | Annotation |  | Assembly |  | Annotation |  |
|  | Proteins | Percentage (%) | Proteins | Percentage (%) | Proteins | Percentage (%) | Proteins | Percentage (%) |
| Complete BUSCOs | 7,191 | 99.78 | 7,158 | 99.32 | 7,189 | 99.75 | 7,169 | 99.47 |
| Complete Single-Copy BUSCOs | 7,170 | 99.49 | 7,135 | 99 | 7,168 | 99.46 | 7,145 | 99.14 |
| Complete Duplicated BUSCOs | 21 | 0.29 | 23 | 0.32 | 21 | 0.29 | 24 | 0.33 |
| Fragmented BUSCOs | 1 | 0.01 | 8 | 0.11 | 2 | 0.03 | 8 | 0.11 |
| Missing BUSCOs | 15 | 0.21 | 41 | 0.57 | 16 | 0.22 | 30 | 0.42 |
| Total BUSCO groups searched | 7,207 | 100 | 7,207 | 100 | 7,207 | 100 | 7,207 | 100 |
| Database | actinopterygii_odb12 |  |  |  | actinopterygii_odb12 |  |  |  |

**Table 14.** Consensus quality assessment.

| reads_type | Yushu_Calti_1.0 |  | Yushu_Efusc_1.0 |  |
| --- | --- | --- | --- | --- |
|  | short read data | PacBio long read data | short read data | PacBio long read data |
| K_asm | 505,574 | 36,245 | 535,051 | 26,009 |
| K_total | 1,077,333,796 | 1,077,333,796 | 1,083,364,142 | 1,083,364,142 |
| QV | 46.295 | 57.7413 | 46.0731 | 59.2068 |
| Error rate | 2.35E-05 | 1.68E-06 | 2.47E-05 | 1.20E-06 |

## Code availability

No custom code was used in the processing of the datasets described in this study. All bioinformatics software and pipelines were run according to the developers’ instructions and protocols. The software versions and specific parameters used are listed in the Methods section.

## Acknowledgements

This work was supported by Specific Research Fund of The Innovation Platform for Academicians of Hainan Province (YSPTZX202402) and Key Research and Development Project of Shandong Province (2024CXPT071-1).

## Author contributions

S.C. conceived the project and revised the manuscript. Y.C. analyzed data and drafted the manuscript. C.W. & S.L. supervised the data and revised the manuscript. M.L., Y.Y. & C.C. collected and prepared the sequencing samples.

## Competing interests

The authors declare that they have no known competing interests or personal relationships that could have appeared to influence the work reported in this paper.

## Notes

https://identifiers.org/ncbi/insdc.sra:SRR40183358

https://identifiers.org/ncbi/insdc.sra:SRR40183353

https://identifiers.org/ncbi/insdc.sra:SRR40183354

https://identifiers.org/ncbi/insdc.sra:SRR40183355

https://identifiers.org/ncbi/insdc.sra:SRR40183352

https://identifiers.org/ncbi/insdc:JCDUVQ000000000

https://identifiers.org/ncbi/insdc:JCDUVR000000000

https://doi.org/10.6084/m9.figshare.33264639

### Competing Interest Statement

The authors have declared no competing interest.

## References

1 Ding, S., Liu, Q., Wu, H. & Qu, M. A review of research advances on the biology and artificial breeding of groupers. Journal of Fishery Sciences of China 25, 737 (2018).

2 Liu, M. & de Mitcheson, Y. S. Gonad development during sexual differentiation in hatchery-produced orange-spotted grouper (Epinephelus coioides) and humpback grouper (Cromileptes altivelis) (Pisces: Serranidae, Epinephelinae). Aquaculture 287, 191–202 (2009).

3 Mo, Z. et al. Transcriptomic analysis reveals innate immune mechanisms of an underlying parasite-resistant grouper hybrid (Epinephelus fuscogutatus × Epinephelus lanceolatus). Fish & Shellfish Immunology 119, 67–75 (2021).

4 Sun, Y. et al. Transcriptome analysis reveals the molecular mechanisms underlying growth superiority in a novel grouper hybrid (Epinephelus fuscogutatus♀ × E. lanceolatus♂). BMC Genetics 17 (2016).

5 Chen, S. et al. Heterosis in growth and low temperature tolerance in Jinhu grouper (Epinephelus fuscoguttatus ♀ × Epinephelus tukula ♂). Aquaculture 562 (2023).

6 Li, H. & Durbin, R. Genome assembly in the telomere-to-telomere era. Nature Reviews Genetics 25, 658–670 (2024).

7 Nurk, S. et al. The complete sequence of a human genome. Science 376, 44–53 (2022).

8 Ge, H. et al. De novo assembly of a chromosome-level reference genome of red-spotted grouper (Epinephelus akaara) using nanopore sequencing and Hi-C. Molecular Ecology Resources 19, 1461–1469 (2019).

9 Han, W. et al. Improved chromosomal-level genome assembly and re-annotation of leopard coral grouper. Scientific Data 10 (2023).

10 Zhou, Q., Gao, H., Xu, H., Lin, H. & Chen, S. A Chromosomal-scale Reference Genome of the Kelp Grouper Epinephelus moara. Marine Biotechnology 23, 12–16 (2020).

11 Zeng, L. et al. Chromosome-level genome assembly and annotation of potato grouper (Epinephelus tukula). Scientific Data 13 (2026).

12 Yang, Y. et al. Whole-genome sequencing of brown-marbled grouper (Epinephelus fuscoguttatus) provides insights into adaptive evolution and growth differences. Molecular Ecology Resources 22, 711–723 (2021).

13 Yang, Y. et al. Chromosome Genome Assembly of Cromileptes altivelis Reveals Loss of Genome Fragment in Cromileptes Compared with Epinephelus Species. Genes 12 (2021).

14 Liu, J. et al. Chromosome-level genome assembly of humpback grouper using PacBio HiFi reads and Hi-C technologies. Scientific Data 11 (2024).

15 Khalifa, R. et al. Chromosome-level genome assembly of the hamour (orange-spotted grouper), Epinephelus coioides. F1000Research 14 (2025).

16 Liu, Y. & Mao, Y. Chromosome-Level Genome Assembly and Genomic Analysis of the Hybrid Grouper ShanHu (Epinephelus fuscoguttatus ♀ × Epinephelus polyphekadion ♂). International Journal of Molecular Sciences 26 (2025).

17 Zhou, Q. et al. Telomere-to-telomere gapless genome assembly of the giant grouper (Epinephelus lanceolatus). Scientific Data 11 (2024).

18 Lu, S. et al. Gap-free telomere-to-telomere haplotype assembly of the tomato hind (Cephalopholis sonnerati). Scientific Data 11, 1268 (2024).

19 Wang, L. et al. A telomere-to-telomere genome assembly and annotation of Epinephelus fasciatus (the blacktip grouper). Scientific Data (2026).

20 Zhao, B. et al. A Complete Telomere-To-Telomere Assembly of Plectropomus leopardus and Phylogenomic Insights Into Perciformes. Evolutionary Applications 19 (2026).

21 Chen, S., Zhou, Y., Chen, Y. & Gu, J. fastp: an ultra-fast all-in-one FASTQ preprocessor. Bioinformatics 34, i884–i890 (2018).

22 Andrews, S. FastQC: a quality control tool for high throughput sequence data. Babraham Bioinformatics (2010).

23 Altschul, S. F., Gish, W., Miller, W., Myers, E. W. & Lipman, D. J. Basic local alignment search tool. Journal of Molecular Biology 215, 403–410 (1990).

24 Marçais, G. & Kingsford, C. A fast, lock-free approach for efficient parallel counting of occurrences of k-mers. Bioinformatics 27, 764–770 (2011).

25 Liu, B. et al. Estimation of genomic characteristics by analyzing k-mer frequency in de novo genome projects. Quantitative Biology 1, 62–67 (2013).

26 Ranallo-Benavidez, T. R., Jaron, K. S. & Schatz, M. C. GenomeScope 2.0 and Smudgeplot for reference-free profiling of polyploid genomes. Nature Communications 11, 1432 (2020).

27 Cheng, H., Concepcion, G. T., Feng, X., Zhang, H. & Li, H. Haplotype-resolved de novo assembly using phased assembly graphs with hifiasm. Nature Methods 18, 170–175 (2021).

28 Roach, M. J., Schmidt, S. A. & Borneman, A. R. Purge Haplotigs: allelic contig reassignment for third-gen diploid genome assemblies. BMC Bioinformatics 19, 460. (2018).

29 Wingett, S. W. et al. HiCUP: pipeline for mapping and processing Hi-C data. F1000Research 4, 1310 (2015).

30 Durand, N. C. et al. Juicer Provides a One-Click System for Analyzing Loop-Resolution Hi-C Experiments. Cell Systems 3, 95–98 (2016).

31 Olga Dudchenko, Sanjit S. Batra, Arina D. Omer & Nyquist, S. K. De novo assembly of the Aedes aegypti genome using Hi-C yields chromosome-length scaffolds. Science 356, 92–95 (2017).

32 Zeng, X. et al. Chromosome-level scaffolding of haplotype-resolved assemblies using Hi-C data without reference genomes. Nature Plants 10, 1184–1200 (2024).

33 Durand, N. C. et al. Juicebox Provides a Visualization System for Hi-C Contact Maps with Unlimited Zoom. Cell Systems 3, 99–101 (2016).

34 Xu, M. et al. TGS-GapCloser: A fast and accurate gap closer for large genomes with low coverage of error-prone long reads. GigaScience 9, giaa094. (2020).

35 Benson, G. Tandem repeats finder: a program to analyze DNA sequences. Nucleic Acids Research 27, 573–580 (1999).

36 Smit, A. F. Interspersed repeats and other mementos of transposable elements in mammalian genomes. Current Opinion in Genetics & Development 9, 657–663 (1999).

37 Price AL, Jones NC & Pa, P. De novo identification of repeat families in large genomes. Bioinformatics 21, i351–i358 (2005).

38 Ou, S. & Jiang, N. LTR_FINDER_parallel: parallelization of LTR_FINDER enabling rapid identification of long terminal repeat retrotransposons. Mobile DNA 10 (2019).

39 Slater, G. S. C. & Birney, E. Automated generation of heuristics for biological sequence comparison. BMC Bioinformatics 6 (2005).

40 Shumate, A. & Salzberg, S. L. Liftoff: accurate mapping of gene annotations. Bioinformatics 37, 1639–1643 (2021).

41 Stanke, M. et al. AUGUSTUS: ab initio prediction of alternative transcripts. Nucleic Acids Research 34, W435–W439 (2006).

42 Burge, C. & Karlin, S. Prediction of complete gene structures in human genomic DNA. Journal of Molecular Biology 268, 78–94 (1997).

43 Kim, D., Paggi, J. M., Park, C., Bennett, C. & Salzberg, S. L. Graph-based genome alignment and genotyping with HISAT2 and HISAT-genotype. Nature Biotechnology 37, 907–915 (2019).

44 Pertea, M. et al. StringTie enables improved reconstruction of a transcriptome from RNA-seq reads. Nature Biotechnology 33, 290–295 (2015).

45 Li, H. Minimap2: pairwise alignment for nucleotide sequences. Bioinformatics 34, 3094–3100 (2018).

46 Huang, N. & Li, H. compleasm: a faster and more accurate reimplementation of BUSCO. Bioinformatics 39, btad595. (2023).

47 Holt, C. & Yandell, M. MAKER2: an annotation pipeline and genome-database management tool for second-generation genome projects. BMC Bioinformatics 12 (2011).

48 Buchfink, B., Reuter, K. & Drost, H.-G. Sensitive protein alignments at tree-of-life scale using DIAMOND. Nature Methods 18, 366–368 (2021).

49 Sayers, E. W. et al. Database resources of the national center for biotechnology information. Nucleic Acids Research 50, D20–D26 (2022).

50 Coudert, E. et al. Annotation of biologically relevant ligands in UniProtKB using ChEBI. Bioinformatics 39, btac793 (2023).

51 Tatusov, R. L. et al. The COG database: an updated version includes eukaryotes. BMC Bioinformatics 4, 41 (2003).

52 Roberts, R. J., Vincze, T., Posfai, J. & Macelis, D. REBASE: a database for DNA restriction and modification: enzymes, genes and genomes. Nucleic Acids Research 51, D629–D630 (2023).

53 Fischer, M. et al. The Cytochrome P450 Engineering Database: a navigation and prediction tool for the cytochrome P450 protein family. Bioinformatics 23, 2015–2017 (2007).

54 Saier, M. H. et al. The Transporter Classification Database (TCDB): 2021 update. Nucleic Acids Research 49, D461–D467 (2021).

55 Shen, W.-K. et al. AnimalTFDB 4.0: a comprehensive animal transcription factor database updated with variation and expression annotations. Nucleic Acids Research 51, D39–D45 (2023).

56 Aramaki, T. et al. KofamKOALA: KEGG Ortholog assignment based on profile HMM and adaptive score threshold. Bioinformatics 36, 2251–2252 (2020).

57 Kanehisa, M. & Goto, S. KEGG: Kyoto Encyclopedia of Genes and Genomes. Nucleic Acids Research 28, 27–30 (2000).

58 Paysan-Lafosse, T. et al. InterPro in 2022. Nucleic Acids Research 51, D418–D427 (2023).

59 Ashburner, M. et al. Gene Ontology: tool for the unification of biology. Nature Genetics 25, 25–29 (2000).

60 Finn, R. D. et al. Pfam: the protein families database. Nucleic Acids Research 42, D222–D230 (2014).

61 Jones, P. et al. InterProScan 5: genome-scale protein function classification. Bioinformatics 30, 1236–1240 (2014).

62 Zhang, H. et al. dbCAN2: a meta server for automated carbohydrate-active enzyme annotation. Nucleic Acids Research 46, W95–W101 (2018).

63 Krogh, A., Larsson, B., von Heijne, G. & Sonnhammer, E. L. L. Predicting transmembrane protein topology with a hidden markov model: application to complete genomes11Edited by F. Cohen. Journal of Molecular Biology 305, 567–580 (2001).

64 Teufel, F. et al. SignalP 6.0 predicts all five types of signal peptides using protein language models. Nature Biotechnology 40, 1023–1025 (2022).

65 Horton, P. et al. WoLF PSORT: protein localization predictor. Nucleic Acids Research 35, W585–W587 (2007).

66 Nawrocki, E. P. & Eddy, S. R. Infernal 1.1: 100-fold faster RNA homology searches. Bioinformatics 29, 2933–2935 (2013).

67 Chan, Patricia P., Lin, Brian Y., Mak, Allysia J. & Lowe, Todd M. tRNAscan-SE 2.0: improved detection and functional classification of transfer RNA genes. Nucleic Acids Research 49, 9077–9096 (2021).

68 NCBI Sequence Read Archive: SRR40183358 https://identifiers.org/ncbi/insdc.sra:SRR40183358 (2026).

69 NCBI Sequence Read Archive: SRR40183353 https://identifiers.org/ncbi/insdc.sra:SRR40183353 (2026).

70 NCBI Sequence Read Archive: SRR40183354 https://identifiers.org/ncbi/insdc.sra:SRR40183354 (2026).

71 NCBI Sequence Read Archive: SRR40183355 https://identifiers.org/ncbi/insdc.sra:SRR40183355 (2026).

72 NCBI Sequence Read Archive: SRR40183352 https://identifiers.org/ncbi/insdc.sra:SRR40183352 (2026).

73 NCBI GenBank: JCDUVQ000000000 https://identifiers.org/ncbi/insdc:JCDUVQ000000000 (2026).

74 NCBI GenBank: JCDUVR000000000 https://identifiers.org/ncbi/insdc:JCDUVR000000000 (2026).

75 Youfeng, C. Genome sequencing and assembly of the hybrid grouper “Yushuban” (Cromileptes altivelis × Epinephelus fuscoguttatus). 10.6084/m9.figshare.33264639 (2026).

76 Li, H. & Durbin, R. Fast and accurate short read alignment with Burrows–Wheeler transform. Bioinformatics 25, 1754–1760 (2009).

77 Manni, M., Berkeley, M. R., Seppey, M., Simão, F. A. & Zdobnov, E. M. BUSCO Update: Novel and Streamlined Workflows along with Broader and Deeper Phylogenetic Coverage for Scoring of Eukaryotic, Prokaryotic, and Viral Genomes. Molecular Biology and Evolution 38, 4647–4654 (2021).

78 Rhie, A., Walenz, B. P., Koren, S. & Phillippy, A. M. Merqury: reference-free quality, completeness, and phasing assessment for genome assemblies. Genome Biology 21, 245. (2020).

