## supplemental table S1, Figure S1 and Figure S2 for "Gap-free telomere-to-telomere haplotypes assembly of the hybrid grouper ‘Yushuban’ (*Cromileptes altivelis* ♀ × *Epinephelus fuscoguttatus* ♂)"

| Content | Page |
| --- | --- |
| Table S1 Next-generation resequencing data from fin-clip samples of non-parental individuals of the two parental species (*C. altivelis* and *E. fuscoguttatus*). | 1 |
| Fig. S1 Circular plot of sequencing depth distribution from next-generation sequencing data. | 2 |
| Fig. S2 Genomic collinearity between the two haplotypes of *C. altivelis* × *E. fuscoguttatus*. | 3 |

| Sample Name | FB-F (Female parent) | MB-F (Male parent) |
|---|---|---|
| parental species | *Cromileptes altivelis* | *Epinephelus fuscoguttatus* |
| Total Raw Reads(M) | 332.41 | 356.07 |
| Total Clean Reads(M) | 331.97 | 355.78 |
| Total Raw Bases(Gb) | 49.86 | 53.41 |
| Total Clean Bases(Gb) | 49.28 | 52.94 |
| Clean Reads Ratio(%) | 99.87 | 99.92 |
| Clean Bases Ratio(%) | 98.84 | 99.13 |
| Q20(%) | 99.19 | 99.18 |
| Q30(%) | 97.89 | 97.9 |
| GC_Content(%) | 42.07 | 41.59 |

Table S1 Next-generation resequencing data from fin-clip samples of non-parental individuals of the two parental species (*C. altivelis* and *E. fuscoguttatus*).

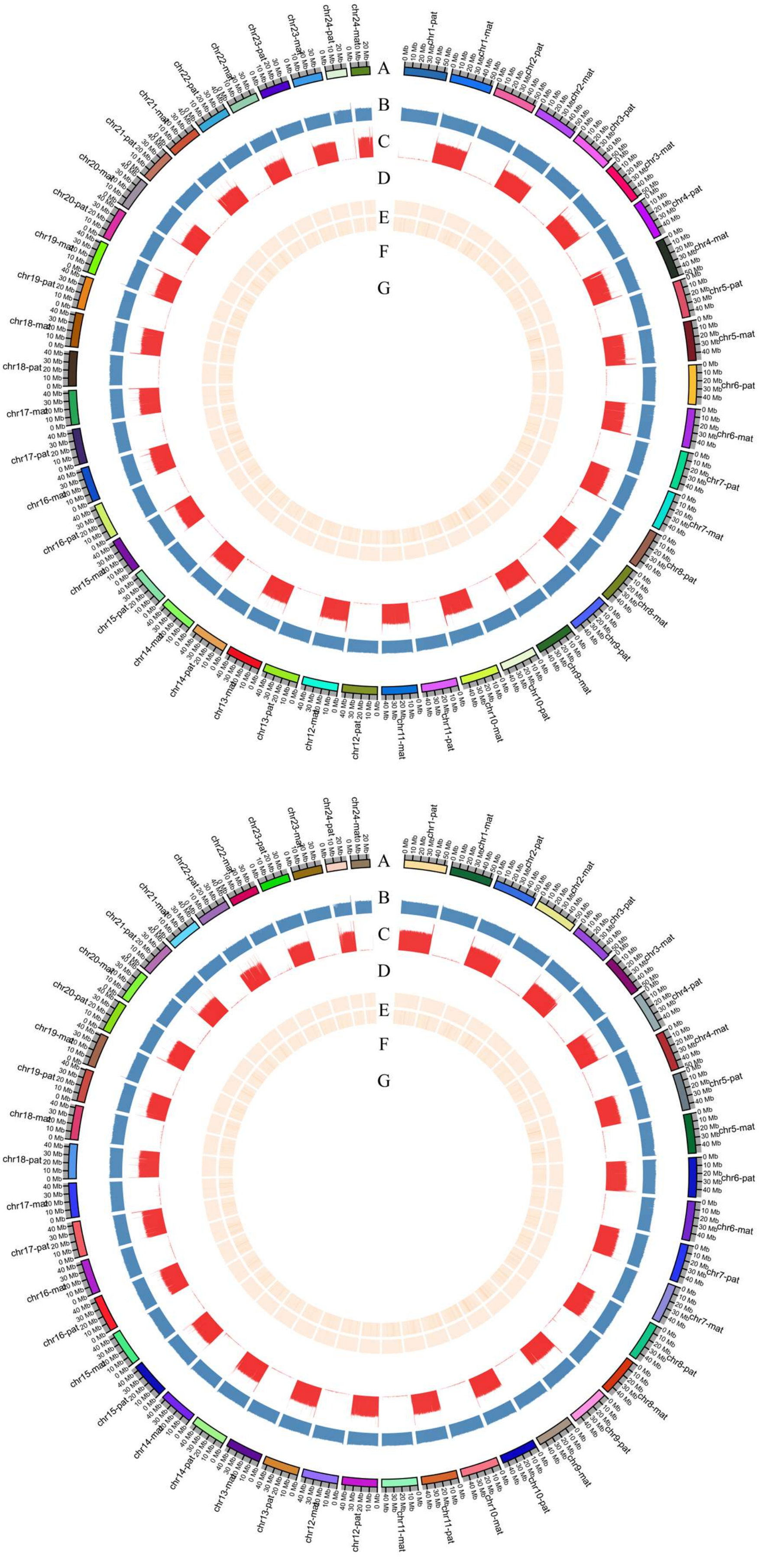


Fig. S1 Circular plot of sequencing depth distribution from next-generation sequencing data. Hap2(upper panel): aligned to female parent *C. altivelis*; Hap1(lower panel): aligned to male parent, *E. fuscoguttatus*. Outer to inner rings: chromosome labels, GC content, and (c)alignment depth of the paternal parent (non-offspring).

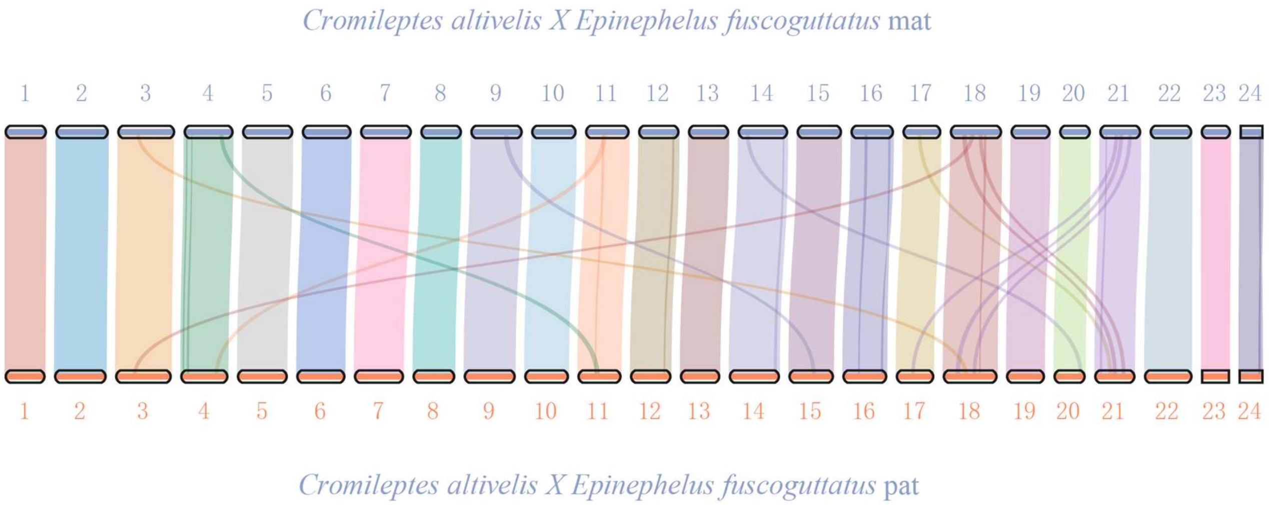


Fig. S2 Genomic collinearity between the two haplotypes of *C. altivelis* × *E. fuscoguttatus*.
